# Transcriptome analysis revealed the possible contribution of chromosome introgression fragments from Sea Island cotton (*Gossypium barbadense*) to fiber strength in Upland cotton (*Gossypium hirsutum*)

**DOI:** 10.1101/073726

**Authors:** Quanwei Lu, Yuzhen Shi, Xianghui Xiao, Pengtao Li, Juwu Gong, Aiying Liu, Haihong Shang, Wankui Gong, Tingting Chen, Qun Ge, Junwen Li, Wei Su, Shaoqi Li, Zhen Zhang, Youlu Yuan, Jinling Huang

## Abstract

Cotton fiber strength is a critical property determining fiber qualities, and determined by the secondary cell wall development. Understanding the mechanism of fiber development will provide a way to improvement of fiber strength. In this study, the introgression lines of upland and sea island cotton, and have experience of four generations of backcross with upland parent, and have significant higher fiber strength than their upland parent, and the transcriptome were analyzed and compared between the introgression lines and their upland parent. There were 2201 differentially expressed genes (DEG) identified by comparing two introgression lines with their recurrent parent CCRI45, in different development stages from 15 days post-anthesis (DPA) to 28 DPA. The up-regulated genes regulated the polysaccharide metabolic process, single-organism localization, cell wall organization or biogenesis and so on. The down-regulated genes involved in the microtubule-based process, cellular response to stress, cell cycle process and so on. Further functional analysis revealed three significant functional genes, XLOC_036333 (mannosyl-oligosaccharide-alpha-mannosidase mns1), XLOC_029945 (*FLA8*) and XLOC_075372 (*snakin-1*), playing important roles in the regulation of cotton fiber strength. Our results provide important candidates genes and inspirations for the future investigation of the molecular mechanism of fiber quality formation, and improvement of cotton fiber quality in breeding.

## Introduction

Cotton is a world's leading commercial fiber crop, providing us the most prevalent renewable natural fibers used for the global textile industry [1]. Both high fiber quality and high yield in concomitant has become a major goal of current cotton breeding programs [2]. Upland cotton produce high fiber yield with relatively low quality, Sea Island cotton produce high quality fiber with relatively low yield. Upland cotton are commonly crossed with Sea Island cotton for obtain introgression lines, which could produce longer, finer and stronger fiber as well as high yield [3, 4]. The cotton fibers develop from single epidermal cells of the developing ovule, serving as an excellent model for the biologically studies of plant cell elongation, as well as biosynthesis of cell wall and cellulose. After initiation (−3 to +3 days post-anthesis, DPA), the fiber cell elongates in rapid rate with peak elongation more than 2 millimeter per day in *Gossypium. hirsutum* (*G. hirsutum*) (3–23 DPA). Overlapping with the elongation stage, fiber cells synchronously enter the stage of secondary wall deposition (15–40 DPA) and then maturation (40–50 DPA) [5].

Fiber strength is one of an important property of cotton fiber quality, and mainly determined by the secondary cell wall biosynthesis. Cellulose is as the major compontent of fiber (over 90%), it’s synthesis increases by over 100-fold during the secondary cell wall thickening, thereby providing mechanical support and protection to the plant body and fiber strength [6]. Rapid and dynamic changes in genes during this stage have been revealed with a complex involving processes such as transcriptional regulation, signal transduction and metabolic pathways, may contribute to cotton fiber development [7, 8]. Fiber strength is a multigenic trait controlled by number of genes [9]. A few genes involved in secondary cell wall and fiber strength have been identified. For example, *GbEXPA2* overexpression in *G. hirsutum* produced fibers with a slightly thicker wall and increased crystalline cellulose content, and *GbEXPATR* overexpression resulted in longer, finer and stronger fibers coupled with significantly thinner cell wall [10]. In addition, secondary cell wall biosynthesis, hormone responses may also associate with cell elongation [11]. A recent study suggested that receptor-like kinases signaling pathways might regulate cotton fiber cell wall assembly and strength by mediating a coordination of cell elongation and secondary cell wall biosynthesis [11]. Despite these, molecular mechanisms responsible for managing individual fiber strength and secondary wall formation in cotton remain not fully clear. Further investigations are under great requirement.

Genome-wide transcriptome profiling provides a powerful approach for effectively discovering significant candidate genes and pathways, uncovering the genetic and molecular basis of fiber quality traits [7, 12]. Several researches applying transcriptome analysis have provided some insights into the dynamic programming associated with developmental processes [13-15]. Recently, comparative transcriptome analysis focused on fiber development suggested candidate genes for regulating cotton fiber cell wall assembly and strength in *G. hirsutum* line MD52ne [11]. Another recent study of fiber indicated a set of candidate genes for molecular breeding and genetic manipulation of lint yield in cotton [16]. However, there is little study focused on the introgression lines with high-strength fiber (*G. hirsutum* CCRI45) crossed with Sea Island cotton *G. barbadense* Hai1.

In this study, we performed transcriptome sequencing to analyze common fiber strength related genes and pathways in developing fibers. Through comparing upland parent with the introgression lines that developed in our laboratory[17], to understand the molecular mechanisms underlying cotton fiber strength regulation and the formation of superior quality fibers. The results of our study provide new insights for further research to the improvement of breeding.

## Material and method

### Plant material

Upland cotton *G. hirsutum* CCRI45 and Sea Island cotton *G. barbadense* Hai1 were crossed to construct backcross populations. The detailed method of developing chromosome segment introgression lines has been described previously[17]. There were four kinds of cotton: CCRI45, MBI7561, MBI7747 and MBI7285. (Table 1).

**Table 1.** The average performance of primary fiber yield and quality traits for CCRI45 and Hai1.

| Cotton | Boll weight (g) | Ginning outturn (%) | Upper Half Mean Length (mm) | Mrc | Fiber strength (cN/tex) |
| --- | --- | --- | --- | --- | --- |
| Hai1 | 3.20 | 32.50 | 32.93 | 4.6 | 42.37 |
| MBI7561 | 6.38 | 41.86 | 32.10 | 5.10 | 34.30 |
| MBI7747 | 5.75 | 35.95 | 33.80 | 4.30 | 36.50 |
| MBI7285 | 4.35 | 36.61 | 27.10 | 4.40 | 25.90 |
| CCRI45 | 5.38 | 35.24 | 28.13 | 5.03 | 27.97 |

The fiber samples were collected at 15, 20, 25 and 28 DPA, frozen in liquid nitrogen and stored at −70°C.

### Transcriptome sequencing

Total RNA was extracted from frozen tissue using an improved CTAB extraction protocol. Sequencing libraries were generated using NEBNext^®^ Ultra™ RNA Library rep Kit for Illumina^®^ (NEB, USA) following manufacturer’s recommendations and index codes were added to attribute sequences to each sample.Transcriptome sequencing was performed.

### Data processing, statistical evaluation, and selection of differentially expressed genes(DEGs)

The library fragments were purified with AMPure XP system (Beckman Coulter, Beverly, USA). cDNA fragments of preferentially 150∼200 bp in length were selected A database of 150∼200 bp sequences was produced beginning with CATG using 66,434 reference genes from the allotetraploid species *G. hirsutum* (http://mascotton.njau.edu.cn). The remaining high quality sequences were then mapped to this database; only a single mismatch was allowed, and more than one match was excluded. HTSeq v0.6.1 was used to count the reads numbers mapped to each gene.[18] Expression levels are expressed as RPKM, transcripts per million. To identify DEGs during fiber development (secondary cell wall formation stage), we compared pairs of DEG profiles using the GFOLD method [19] from different libraries. The genes with |GFOLD value|>1 acted as DEGs.

### Cluster analysis and function analysis

DEG clustering in introgression lines at different developmental stages were performed with the ‘Self-organizing tree algorithm’ (SOTA, Multiple Array Viewer software, MeV 4.9.0). GO enrichment was done using BLAST2GO (http://www.blast2go.com/b2ghome).

### Quantitative real-time PCR (qRT-PCR)

Real-time PCR of the selected differentially expressed genes was performed using 7500 Fast (Applied Biosystems, San Francisco, CA, USA). The gene specific primer were designed by software Primer Premier 5.0, checked by Blastn, and synthesized by the manufacturer (Life Technologies). All reactions were performed in three biological and technological replicates. The cotton histone3 gene were used as internal reference. (F: 5’-GGTGGTGTGAAGAAGCCTCAT-3’, and R:5’-AATTTCACGAACAAGCCTCTGGAA-3’). The relative gene expression levels were presented as 2-ΔCT. The specific primer sequences were listed in Table S1.

## Results

### The phenotypes of four germplasms

In this study, the fiber strength of CCRI45 and the BC_4_F_3:5_ individuals MBI7561, MBI7747 and MBI7285 was 27.97 cN/tex,34.30 cN/tex,36.50 cN/tex and 25.90 cN/tex, respectively (Table 1). The upper-half mean length of CCRI45 and the BC4F3:5 individuals MBI7561, MBI7747 and MBI7285 was 28.13 mm,32.10 mm,33.80 and 27.10, respectively (Table 1). The mean strength of MBI7561 and MBI7747 were higher than that of CCRI45, while the mean strength of MBI7285 were lower than that of CCRI45. These CSSLs were used as for the study of functional genetic study of cotton fiber trait.

### Gene expression patterns during cotton fiber development

In order to systematically analyze transcriptome profiles related to cotton fiber development, we performed RNA sequencing for fibers (15, 20, 25and 28 DPA) in introgression lines of MBI7747, MBI7561, MBI7285 and their recurrent parent CCRI45. A total of 44,867 genes were identified from all the libraries. To examine the relationship between the experimental samples, Pearson correlation coefficient (PCC) analysis was performed on all the genes obtained from all libraries. As shown in Fig 1a, the gene expression patterns in CCRI45 had low similarities at all four stages of fiber development, and low similarities were also found in all libraries at the early stages of 15 DPA. However, higher similarities were observed at later stages (20 DPA, 25 DPA and 28 DPA) compared to the earlier stages (Fig 1a). These findings revealed that the gene expression patterns altered more remarkably in MBI7561, MBI7747 and MBI7285 in the later stages than in the earlier stages, and these results may because the three kinds of cotton carried distinct *G. barbadense* chromosomal segments.

**Fig 1.**
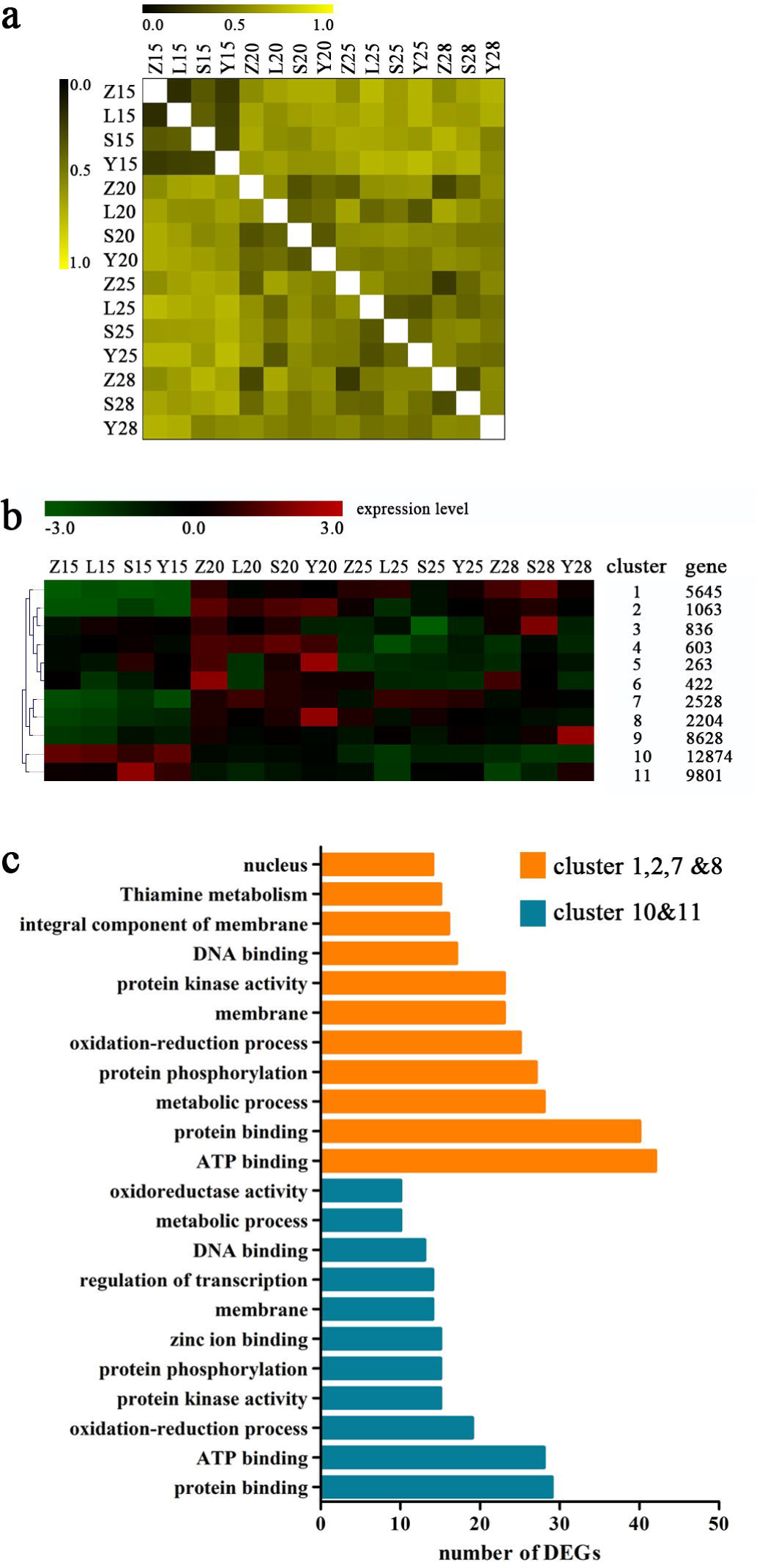
Statically analysis of transcript profiling data. Z, S, L and Y indicated CCRI45, MBI7561, MBI7747 and MBI7285 respectively. 15, 15 DPA; 20, 20 DPA; 25, 25 DPA; 28, 28 DPA. (a) Pearson correlation coefficient analysis of all samples. Z, S, L and Y indicated CCRI45, MBI7561, MBI7747 and MBI7285 respectively. 15, 15 DPA; 20, 20 DPA; 25, 25 DPA; 28, 28 DPA. (b) SOTA clustering analysis of genes in these samples. (c) Function annotation of genes in special clusters.

To further analyze the fiber transcriptomes at different developmental stages, the 44867 genes were classified into 11 groups (Fig 1b). Genes in cluster 10 and 11 were more highly expressed in the 15 DPA than other DPAs, but genes in cluster 1, 2, 7 and 8 showed the opposite expression pattern. Annotation of the gene functions, genes in cluster 1, 2, 7 and 8 highly regulated protein binding and ATP binding, and genes in cluster 10 and 11 also did (Fig 1c). However, the genes in cluster 1, 2, 7 and 8 specially regulated the functions such as integral component of membrane, and thiamine metabolism. The genes of cluster 10 and 11 were mainly related to the regulation of transcription, zinc ion binding, and etc.

### Analysis of differentially expressed genes (DEGs)

We calculated the DEGs in different developmental stages using GFOLD method and 2201 DEGs were acquired. Most of the DEGs were down-regulated (Fig 2a). The genes were significantly differentially expressed between MBI7747 andCCRI45 in the 15, 20 and 25 DPA (Table 2). In the 25 DPA, there were 615 DEGs expressed between MBI7747 and CCRI45, which was much more than MBI7561 *vs* CCRI45 (543) and MBI7285 *vs* CCRI45 (134) (Fig 2b).

**Fig 2.**
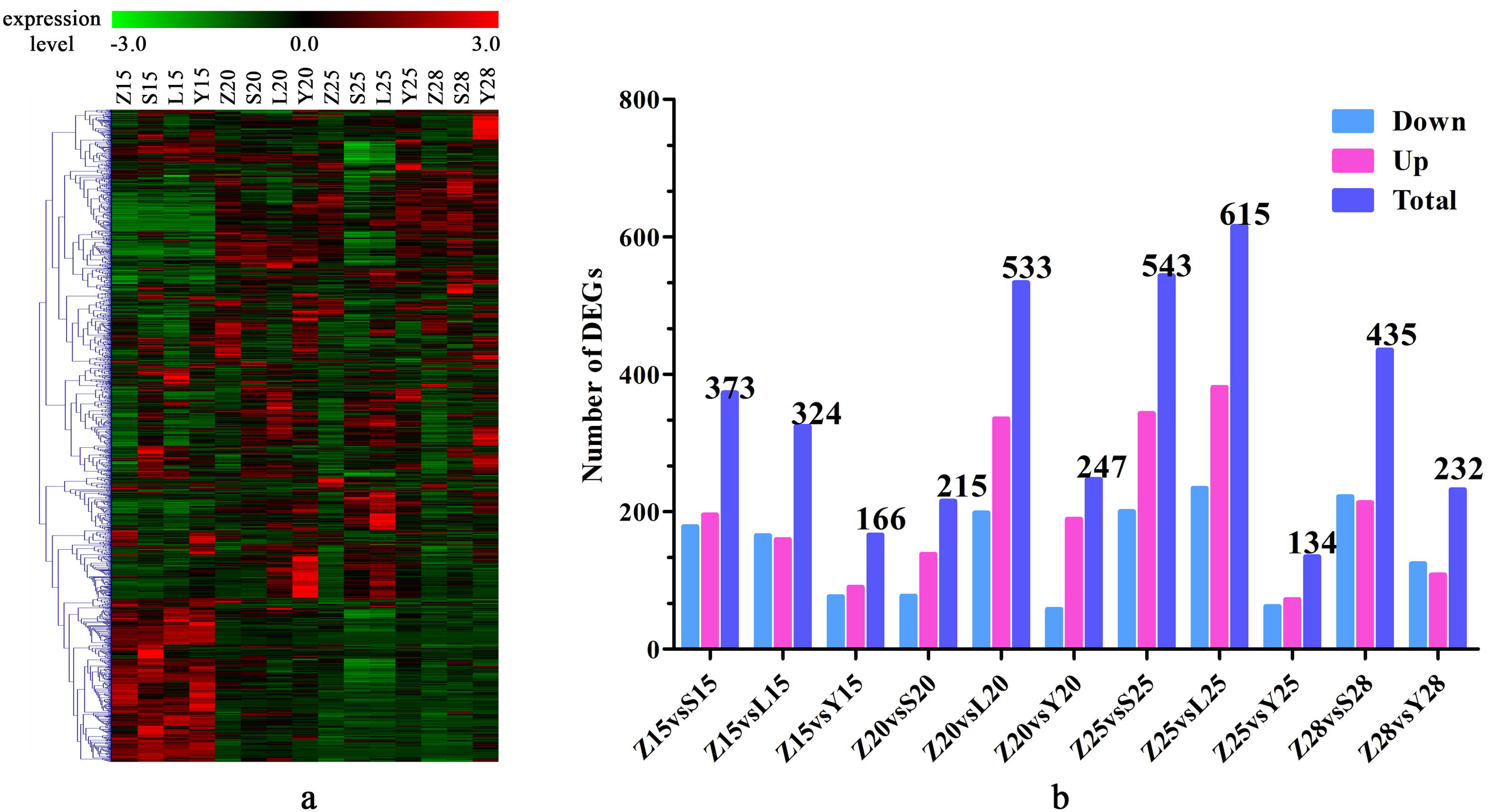
The overview of differentially expressed genes. Z, S, L and Y indicated CCRI45, MBI7561, MBI7747 and MBI7285 respectively. 15, 15 DPA; 20, 20 DPA; 25, 25 DPA; 28, 28 DPA. (a) The heatmap of all the DEGs which were integrated form diverse comparisons. (b) The statistics of DEGs in diverse comparisons.

**Table 2.**
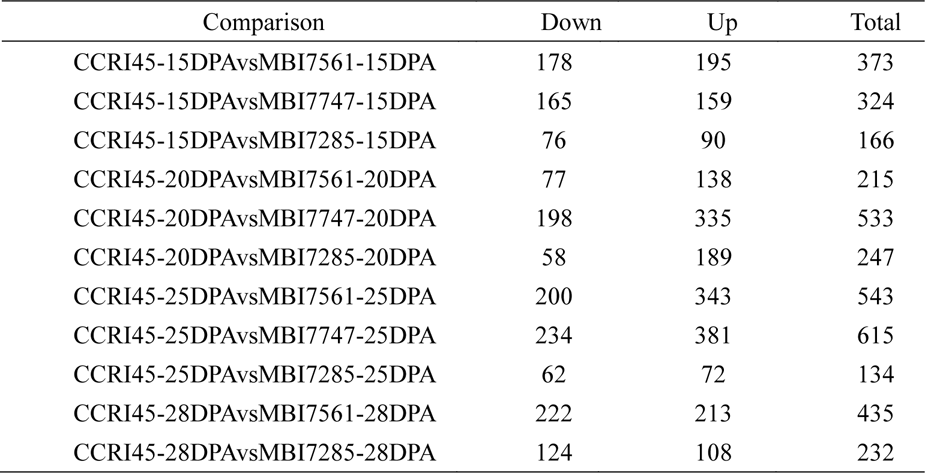
All the DEG in diverse comparison.

Applying the fisher exact test, we enriched the DEGs in biological process. The up-regulated genes regulated the polysaccharide metabolic process, regulation of hormone levels, regulation of transcription and cell wall biogenesis (Fig 3a). The down-regulated genes involved in the microtubule-based process, cellular response to stress, cell cycle process (Fig 3b).

**Fig 3.**
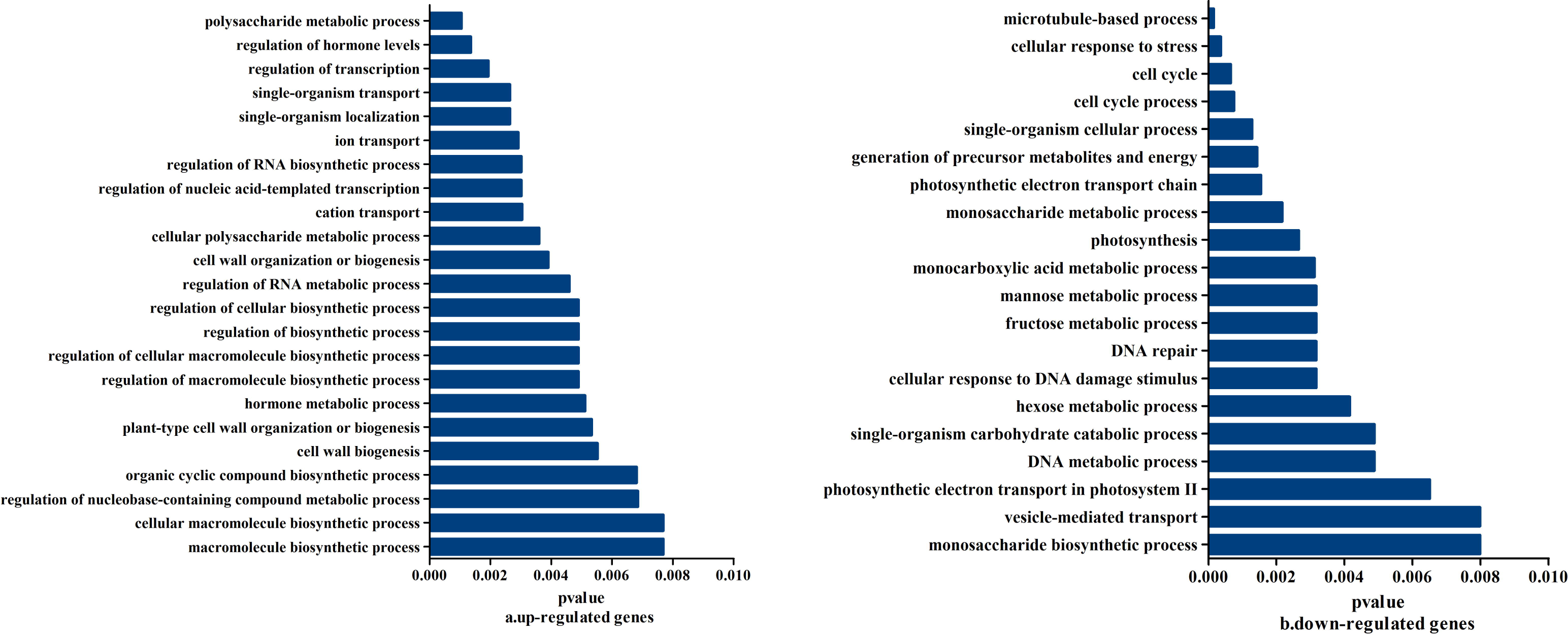
The function enrichment of DEGs. (a). the biological process enriched by the up-regulated genes; (b) the biological process enriched by the down-regulated genes.

### Cluster analysis of DEGs between CCRI45 and MBI7561

To understand the mechanisms underlying the alterations in fiber strength observed in our samples, we analyzed the DEGs among MBI7561 with CCRI45 at different stages, and 19 common DEGs were acquired (Fig 4a). Moreover, we found that the common DEGs were significantly differentially expressed in MBI7561 *vs* CCRI45 and MBI7747 *vs* CCRI45 (Fig 4b). The MBI7561 and MBI7747 samples were the stronger fiber, we examine the expression patterns of all the 756 DEGs of MBI7561 *vs* CCRI45, with cluster analysis method. Finally, 11 clusters were acquired (Fig 4c) and these clusters.

**Fig 4.**
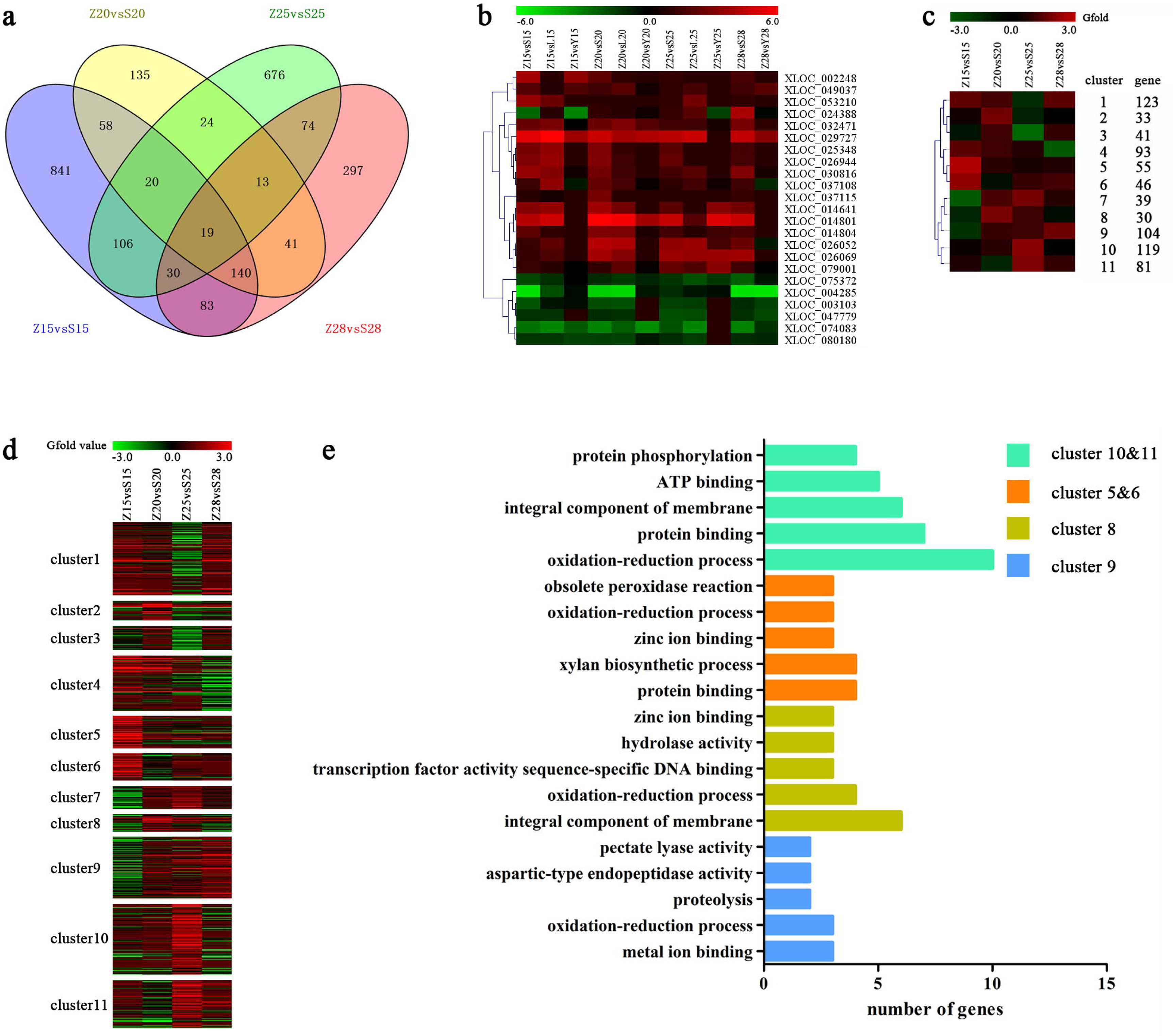
The cluster analysis of DEGs in between CCRI45 and MBI7561. Z, S, L and Y indicated CCRI45, MBI7561, MBI7747 and MBI7285 respectively. 15, 15 DPA; 20, 20 DPA; 25, 25 DPA; 28, 28 DPA. (a) the number of DEGs in diverse comparison. (b) The heatmap of common DEGs. (c) SOTA clustering analysis of all DEGs in diverse comparison between CCRI45 and MBI7561. (d) heatmap analysis of the expression of all DEGs between Z and S. (e) function annotation of DEGs in special clusters.

From the results, the genes in cluster 5 and 6 were highly expressed in 15 DPA; genes in cluster 8 were highly expressed in 20 DPA; genes in cluster10 and 11 were markedly high-expressed in 25 DPA and genes in cluster 9 were highly expressed in 28 DPA. Function annotation and enrichment of these genes were performed, the genes in cluster 5 and 6 regulated protein binding and xylan biosynthetic process. Genes in cluster 8 regulated metal ion binding and oxidation-reduction process. Genes in cluster 10 and 11 regulated oxidation-reduction process and protein binding. Genes in cluster 9 regulated integral component of membrane (Fig 4e).

## Special function biological process

Phytohormones play an important regulatory role in various plant growth and developmental processes. In the present study, numerous genes involved in hormone signal transduction and biosynthesis of auxin, BR, ethylene. Several genes involved in cell wall and fatty acid biosynthesis were differentially expressed at various fiber developmental stages of secondary cell wall deposition. Moreover, genes involved in transcription factors also regulated the fiber strength. We acquired 26 special biological processes including cell wall and carbohydrate metabolic process. The whole expression level of genes in these special biological processes was analyzed by integrating the expression profile (Fig 5a). The heatmap indicated that most genes of these biological processes were lowly expressed. After DEGs number count, we found that most DEGs regulated the cell wall, carbohydrate metabolic process and transcription factor activity (Fig 5b).

**Fig 5.**
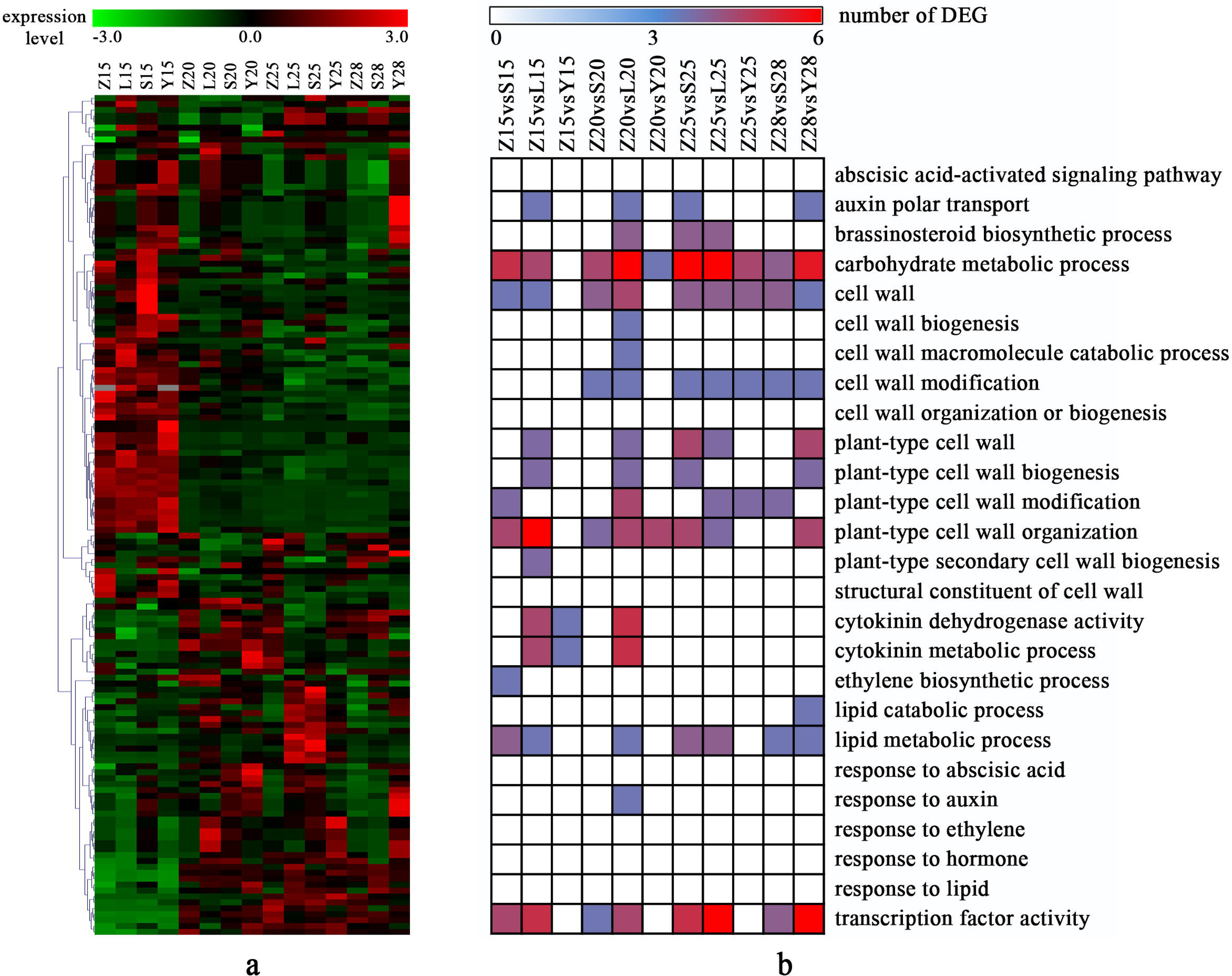
The special biological process. Z, S, L and Y indicated CCRI45, MBI7561, MBI7747 and MBI7285 respectively. 15, 15 DPA; 20, 20 DPA; 25, 25 DPA; 28, 28 DPA. (a) The heatmap of genes in special biological processes. (b) The numbers of DEGs in special biological process.

To study the changes of these biological process in MBI7561 *vs* CCRI45, we analyzed the genes of biological processes in diverse stages which were DEGs in some comparison (Fig 6 and Table S2). As a result, there were three genes expressed in all the four stages: XLOC_036333, XLOC_029945 and XLOC_075372. Expression levels of XLOC_075372 were down-regulated in all the stages. XLOC_075372 was snakin-1, regulated the extracellular region, cell wall and defense response. Expression of XLOC_036333 was up-regulated in 20 DPA, 25 DPA and 28 DPA but not differentially expressed in 15 DPA. XLOC_036333 was mannosyl-oligosaccharide-alpha-mannosidase (*MNS1*), which regulated mannosyl-oligosaccharide 1,2-alpha-mannosidase activity, membrane, calcium ion binding and carbohydrate metabolic process. XLOC_029945 especially up-regulated in 25 DPA comparing with other stages. XLOC_029945 was fasciclin-like arabinogalactan protein 8 (*FLA8*), moderated the plant-type cell wall, anchored component of plasma membrane, regulation of cell size; plant-type cell wall organization, auxin polar transport. These three genes regulated the important biological process and may mediate the secondary cell wall deposition contributing to the development of cotton fiber strength.

**Fig 6.**
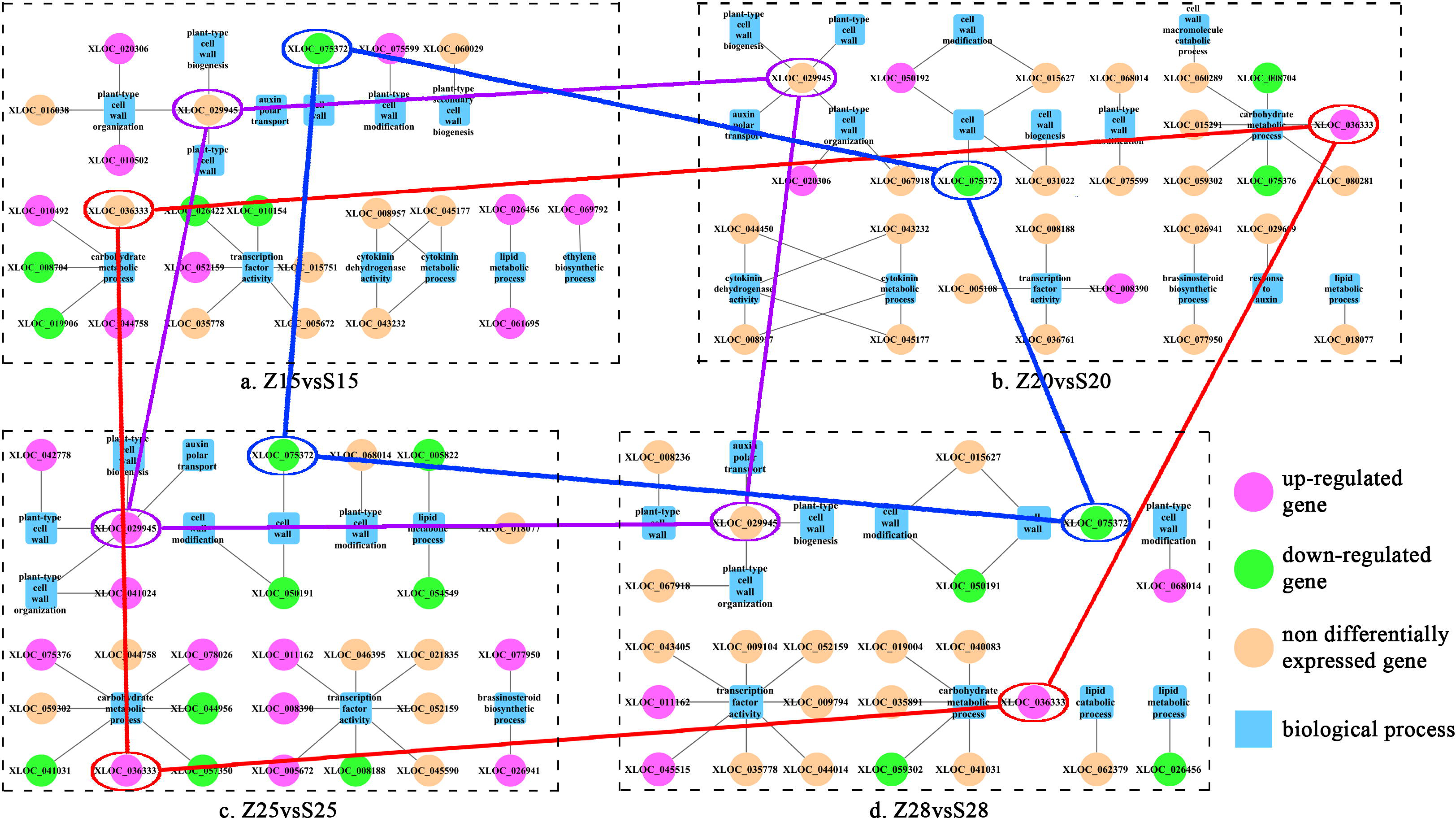
The special biological process and genes in CCRI45 vs MBI7561. Z, S, L and Y indicated CCRI45, MBI7561, MBI7747 and MBI7285 respectively. 15, 15 DPA; 20, 20 DPA; 25, 25 DPA; 28, 28 DPA. (a) the biological processes-genes in 15 DPA. (b) The biological processes-genes in 20 DPA. (c) The biological processes-genes in 25 DPA. (d) The biological processes-genes in 28 DPA. Genes differentially expressed at least one of the 11 comparisons.

To test the reliability of transcriptome data, quantitative RT-PCR analysis was performed to validate the expression of the genes that have been reported in this paper, including XLOC_036333, XLOC_029945 and XLOC_075372 (Fig 7). The expression patterns of these genes were consistent with the results of transcriptome data.

**Fig 7.**
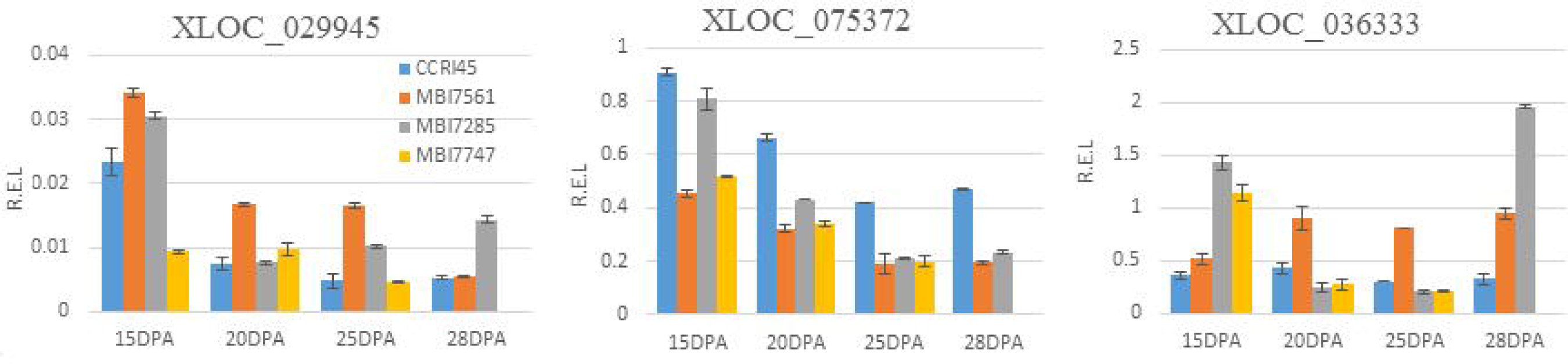
Quantitative RT–PCR validation the expression of the genes that have been reported in this paper. This gene including XLOC_036333, XLOC_029945 and XLOC_075372.

## Discussion

Cotton fiber strength is one of the key traits of fiber quality. It is mainly determined by the secondary cell wall formation stage which affect the yield and quality of fiber, and are important for cotton because of its applications in agriculture. Genetic screens have identified a number of genes involved in fiber development [9]; however, the genes and molecular regulatory mechanisms governing the fiber developmental process and fiber strength are still little understood. In this study, we systematically identified and analyzed cotton fiber strength related gene expression profiles by transcriptome sequencing. We utilized *G. hirsutum* chromosome introgression lines harboring chromosomal segments from *G. barbadense* presenting stronger fiber strength. Through comparing the stronger fiber samples MBI7561 and MBI7747 with their recurrent parent cotton sample CCRI45 at different stages of secondary cell wall formation, 2201 DEGs were obtained and analyzed. Most DEGs were found to regulate the cell wall, carbohydrate metabolic process and transcription factor activity. Moreover, several candidate genes and their involved pathways and biological processes were revealed highly associated with modulating cotton fiber strength. Our results provided some insights for the molecular mechanisms underlying cotton fiber strength regulation and the formation of superior quality fibers.

During recent years, significant progress has been made in large-scale genomic studies, revealing cotton fiber development involved genes and signaling pathways [12, 15, 20, 21]. Transcriptome analysis showed that gene expression patterns and functional distribution were different during cotton fiber development, because that the mechanisms of cotton fiber development are complex with many signal transduction and transcriptional regulation components involved in the process [3, 7, 14]. In our transcriptome analysis, 615 DEGs expressed in MBI7747 *vs* CCRI45 and 543 DEGs in MBI7561 *vs* CCRI45, indicating that gene expression patterns were different in our introgression lines MBI7561 and MBI7747 as comparison with the parent CCRI45. Moreover, DEGs were clustered into groups which altered in different cotton fiber developmental stages. We found cluster 5 and 6 held genes mainly up-regulated at 15 DPA, while genes in cluster 8 up-regulated at 20 DPA, cluster 10 and 11 had increased gene expression at 25 DPA, whereas at 28 DPA genes in cluster 9 were elevated. We also discovered 19 genes consistently differential expression in MBI7561 *vs* CCRI45 and MBI7747 *vs* CCRI45 during all the four DPA stages. Several functional candidates involved in cotton fiber secondary cell wall formation identified, such as polysaccharide metabolic process, regulation of hormone levels, regulation of transcription and cell wall biogenesis. Therefore, our results support the hypothesis that different metabolic pathways and their involved genes can affect fiber strength, and the same pathway in the introgression lines can be altered differentially at various times in development.

In the current study, most of the DEGs were involved in cell wall biogenesis. We proposed that genes implicated with secondary wall synthesis and metabolism should be candidate genes for developing cotton cultivars with superior fiber quality. Actually, mature cotton fiber is composed of nearly pure cellulose (over 90%), and also minor other noncellulosic carbohydrates, such as pectic polysaccharides, xyloglucans, xylans, glucomannans, and glucans [22-24]. Cellulose synthesis in the secondary cell wall of cotton fiber plays a predominant role in fiber cells [24, 25]. Other hemicelluloses and noncellulose are also essential for the normal assembly and mechanical strength of secondary walls. It has been reported that decrease in xylan induces a severe reduction in cellulose deposition, secondary wall thickening and mechanical strength of vessels and fibers [26, 27]. Lignin provides mechanical strength, rigidity and hydrophobicity to secondary walls. A study on *GhMYBL1* transgenic plants discovered enhanced cellulose and lignin biosynthesis [28]. Transcriptional activator *GhMYB7* with potential function upstream of *NAC* transcription factors, may be involved in regulating secondary cell wall biosynthesis of cotton fibers by inducing ectopic deposition of cellulose and lignin [29]. In addition, polysaccharide components such as pectin also play an important role in secondary cell wall synthesis [25]. Higher noncellulosic polysaccharides might trigger a delay in the transition to secondary wall synthesis and responsible for better fiber [10]. It is noteworthy to mention that genes involved in regulation of polysaccharide metabolic process were also up-regulated in our introgression lines.

Furthermore, our results showed that numerous genes involved in hormone signal transduction and biosynthesis of auxin, BR, ethylene. Previous reports revealed that ethylene may promote cell elongation by increasing the expression of sucrose synthase, tubulin, and expansin genes [30]. Activation of ethylene biosynthesis by saturated very-long-chain fatty acids significantly promoted fiber cell elongation in cotton (*G.hirsutum*) [31]. Several DEGs were associated with ethylene biosynthetic process and the interconnected phytohormonal pathways that are involved in superior cotton fiber strength.

Interestingly, we found XLOC_029945 (*FLA8*) up-regulated at 25 DPA in stronger fiber samples. This result revealed that FLA8 may have tissue-specific and fiber secondary cell wall developmental stage-specific expression characters, may contribute to fiber strength by affecting cellulose synthesis and microfibril deposition orientation. *FLA* protein is previously known as a cell-wall-associated protein regulating plant growth and development, such as cell wall pectin plasticizer, cell expansion and cell wall architecture [32]. *FLA* genes were found differentially expressed in different plant tissues and in developing fibers and also differentially between Sea Island cotton and Upland cotton [8, 33, 34]. For example, *GhFLA1* and *GhFLA4* genes highly expressed at 10 DPA whereas *GhFLA2* and *GhFLA6* at 20 DPA contrastively in Upland cotton [35]. Expression level of *GbFL*A5 was significantly higher in Sea Island cotton fibers than in Upland cotton fibers during the secondary cell wall deposition stage from 15 to 45 DPA, and it also played a critical role in regulating fiber strength [33]. These evidences indicated that expression levels of *FLA* genes are diversified and their function in regulating developing cotton fibers may also be variety. These reports also provide supports for our finding that *FLA8* is a potential gene contributing to cotton fiber strength regulation in introgression line. Another gene XLOC_075372 was identified as *snakin-1*. It has been reported that snakin-1 silencing affects cell division, leaf primary metabolism, and cell wall composition in potato plants [36]. Moreover, XLOC_036333 (mannosyl-oligosaccharide-alpha-mannosidase *MNS1*) was up-regulated during 20 DPA to 28 DPA, and was also very potentially related to fiber strength regulation. This gene encodes an enzyme belonged to glycosyl hydrolase family and is involved in the synthesis of glycoproteins. All these three genes may play important roles in the molecular mechanism of cotton fiber strength regulation. However, further investigations are needed for the confirmation.

In conclusion, cotton fiber strength related gene expression profiles were analyzed by comparing the Upland cotton chromosome introgressed lines bearing distinct *G. barbadense* chromosomal segments with the recurrent parent Upland cotton. Different patterns of differentially expressed genes were identified, and altered different metabolic pathways were mainly enriched in secondary cell wall biogenesis, carbohydrate metabolic process and regulation of hormone levels and transcription. Several genes were found to be specifically up-regulated in the stronger fiber cottons at 25 DPA or during 20 DPA to 28 DPA, which may play a significant role in the formation of cotton fiber strength. Our study provides some insights into further research regarding fiber development.

